# DKK1, a negative regulator of Wnt signaling, is a novel inducer of neuroblastoma differentiation

**DOI:** 10.1101/2023.07.25.550446

**Authors:** Shubham Krishna, Bharat Prajapati, Pankaj Seth, Subrata Sinha

## Abstract

**Background:** This study aimed to analyze the role of Dickopff1 [DKK1] as a potential differentiating agent for the neuroblastoma cell line SHSY5Y and neurospheres derived from it.

**Materials and Methods:** SHSY5Y neurospheres were formed from undifferentiated adherent cultures. The cellular properties and gene expression were used to study the effect of DKK1 on SHSY5Y-formed neurospheres. Its effect on SHSY5Y neuronal differentiation was also studied.

**Results:** SHSY5Y adherent undifferentiated cells were grown as neurospheres. Treatment of neurospheres resulted in their fragmentation. It also resulted in reduced mRNA expression of markers of cancer stem cells, pluripotency, and proliferation [p≤0.05]. DKK1 treatment also resulted in reduced mRNA expression of β-catenin and TCF genes. There was significantly higher expression of neuronal differentiation genes in SHSY5Y adherent cells grown in complete DMEM media containing DKK1 as compared to cells grown without DKK1. We also found DKK1 synergized with retinoic acid-induced differentiation of neuroblastoma cells.

**Conclusion:** DKK1 was able to convincingly abrogate neurosphere formation and promote neuronal differentiation of SHSY5Y cells, including synergy with retinoic acid. This was accompanied by corresponding changes in mRNA markers of cancer stem cells, pluripotency and proliferation. These results may have implications for neuroblastoma stemness and differentiation.

## INTRODUCTION

Neuroblastoma is the most common solid childhood cancer originating outside the brain from the primitive sympathetic nervous system, with 90% of cases occurring in children less than 5 years and is responsible for 15% of pediatric cancer death [1,2,3]. Genetic abnormalities in neural crest cells derived sympathoadrenal cells lead to their rapid proliferation, reduced differentiation, and apoptosis of cells [4]. Possible molecules that can be used for neuroblastoma therapy along with chemotherapy include those that promote apoptosis, especially by targeting the p53 and MDM pathway [5], Bcl2 inhibitors [6] electron transport chain inhibitors [7,8] autophagy activators especially by mTOR/AKT inhibitors [9,10,11]. Limited efficacy and the development of resistance, is, however, a feature of these approaches, indicating the need to develop better methods.

Different neuroblastoma-derived cell lines have been used for studying neuroblastoma, of this, one very commonly used cell line is SHSY5Y [12]. It is a subline of the parental line SK-N-SH. SK-N-SH were subcloned to SHSY, then to SHSY5, and finally to SHSY5Y [13]. The SHSY5Y cell line can be differentiated *in vitro* into mature neuronal-like cells by several methods which include treatment with phorbol esters and retinoic acids [14]. One can also lower the SHSY5Y aggregation and increase neurite outgrowth of SHSY5Y cells using dibutyryl cyclic AMP. Retinoic acid has been shown previously used for differentiation therapy in neuroblastoma disease, but has limited efficacy [15]. The MYCN proto-oncogene is an important genetic marker of neuroblastoma tumor aggressiveness. N Myc amplification is seen in aggressive Neuroblastoma [16]. Duffy DJ et al show that retinoic acid and TGF β signaling work together to overcome retinoid resistance. This retinoid resistance is due to MYCN [17]. Bayeva N et al describe how synergistic interactions of retinoic acid derivatives like fenretinide and retinoic acid with molecules like IFN or DHEA have differing effects on neuroblastoma cells [18].

Wnt signaling has been involved in various functions of cell proliferation, stemness, and differentiation. Dysregulation of the Wnt pathway has been linked to the development of various tumors. Wnt-mediated β-catenin activation has both pro and anti-tumor effects in different malignancies There are also differences in the canonical and non-canonical Wnt pathway [19].

Zhang et al show that the Wnt inhibitory factor (WIF-1) functions as a tumor suppressor in SK-N-SH cells by inhibiting the Wnt/β catenin pathway [20]. Szenes M et al have shown that Wnt signaling can drive the proliferation of SK-N-AS cells, but also the differentiation of SK-N-BE(2)-C and SH-SY5Y cells. Active MYCN plays a role Wnt mediated proliferation [21]. The context-dependent differences due to Wnt activation are well known. Becker and Wilting sum up by saying that the canonical and non-canonical roles of Wnt signaling often interact in a mutually inhibitory manner [22].

As reviewed by Becker and Wilting, human neuroblastoma reports show both activation and inhibition of the Wnt pathway with contradictory results [23]. There are reports suggesting inhibition of the Wnt pathway reduces neuroblastoma cell differentiation [24]. Other reports contradict this and suggest that activation of the Wnt pathway is responsible for cancer stemness, leading to chemoresistance. linked proliferation [25,26,27,28]. There is also a report saying inhibition of Wnt signaling promotes neuroblastoma differentiation [29]. To the best of our knowledge, no study shows an in vitro assessment of the Wnt pathway inhibitors on stemness-related pathways in neuroblastoma cells.

Dickkopf -1 or DKK1 is an inhibitor of Wnt signaling. It is well expressed in adrenals and poorly expressed in neuroblastoma tissue and also in neuroblastoma cell lines, like IMR-32, SK-N-SH and SHSY5Y cell lines. In primary neuroblastoma tumor tissue, it has an inverse association with the extent of differentiation [30]. In this study we have used this Wnt inhibitor to study its role in SHSY5Y stemness, proliferation, and differentiation. Neurospheres formed from SHSY5Y cells were treated with DKK 1. This disintegrated the neurosphere, along with a reduction in the expression of stem cell markers. DKK1 also significantly decreases proliferating ability and promotes the neuronal differentiation of SHSY5Y adherent culture, as evident from longer neurites and an increase in neuronal differentiation markers. This effect also synergizes with retinoic acid in increasing neuronal differentiation and reducing proliferation.

## Methods

### Differentiation of SHSY5Y cells using retinoic acid

10% FBS in DMEM was used to grow the undifferentiated cells. For differentiation experiments, the final concentration of 10 μm of retinoic acid was used in complete DMEM media.

### SHSY5Y neurosphere formation

Briefly, in 45.5 ml of neurobasal media, 1ml of Pen-Strep (Invitrogen, USA), 1 ml of NSF1 (Lonza, Charles City, IA), 1 ml of glutamine, 400 μl of N2 supplement, 50 μl of gentamycin, 560 μl of EGF (R&D Systems, Minnesota, United States), 50 μl of FGF (R&D systems, Minnesota, United States), and 0.4 gm of BSA were added and then mixed well. Ultra-low attachment flasks were used for neurosphere formation.

### DKK1 treatment for SHSY5Y

0.5 μg per ml of DKK-1 (Sigma-Aldrich, St. Louis, MO) concentration in DMEM or neurobasal media was used for experiments. For treating neurospheres, cells were grown under appropriate conditions until distinct neurospheres had formed. They were then treated to recombinant DKK1. In adherent SHSY5Y culture, cells were seeded, and after 24 hours, cells were treated to DKK1. After every 24 hours, half of the old media was changed with fresh media containing DKK1 in both neurospheres and adherent SHSY5Y cells.

### RNA isolation and cDNA synthesis

RNA isolation was done according to the established protocol using chloroform - isopropanol method. The cells were scrapped using triazole reagent. Chloroform and isopropanol were added sequentially. The pellet thus formed was washed with ethanol. RNase-free water was then added and heated on the heating block. 1000 ng of RNA were used for cDNA synthesis. cDNA was prepared using a cDNA reverse transcription kit from Applied Biosystems according to their protocol. The cDNA synthesis was done at 25^°^C for 10 minutes, 37^°^C for 2 hours, and 85^°^C for 5 minutes. The sample was then held at 4^°^C.

### Quantitative Real-Time PCR

For quantitative real-time PCR, the cDNA, water, and SYBR green were mixed. It was run on Rotor-Gene Q (Qiagen, Germany). The conditions for amplification were followed: denaturation at 95^°^C for 5 minutes, annealing at 62^°^C for 30 seconds, and amplification at 72^°^C for 1 minute. Amplification was done for 40 to 45 cycles. Primer sequences are given in supplementary S1.

### Immunocytochemistry

The cells were washed with 1X PBS. The cells were fixed in 4% paraformaldehyde in 1 X PBS. The cells were washed with 1X PBS three times for five minutes. We incubated the cells in 4 % BSA and 0.3% triton X 100 in 1X PBS for one hour at room temperature. We incubated the cells in primary antibody in a 0.1% BSA solution at 4^0^C overnight. Cells were washed with 1 X PBS five times for five minutes each. Cells were incubated with a secondary antibody containing 0.1 % BSA for 1 hour at room temperature. Cells were washed with 1 X PBS five times for five minutes each. Cells were once rinsed with milli-Q water. Then the slide was air-dried for 5–7 minutes. Finally, the slide was mounted with an anti-fade reagent containing DAPI. The chamber slide was left at room temperature overnight. The next day, imaging was taken on an apotome.

### Bright-field Imaging

The cells were imaged on a Nikon bright field microscope at 10 X.

### Statistical analysis

The experiments were done in triplicate and were repeated three times. Technical replicates were used for calculating the standard deviation. A student t-test was used to check for a significant difference between the groups. The p-values for significant changes were represented as follows: * p≤0.05; ** p≤0.005; *** p≤0.0005.

## Results

### DKK1 treatment reduces the markers of neuroblastoma stem cells proliferation

SHSY5Y cell line exists in an undifferentiated state (Figures 1A). To study the stemness and pluripotency of SHSY5Y cells, these were grown in a stem cell medium, which resulted in neurosphere formation (Figures 1B). Expression of the stem cell marker nestin was studied in formed neurospheres on PDL-coated slides (Figure 1C), which confirmed the stem cell nature of the neurospheres. DKK1 treatment of the neurospheres for 24 to 48 hours resulted in their fragmentation, with smaller clusters and also some individual cells (Figure 1D-E).

**Figure 1:**
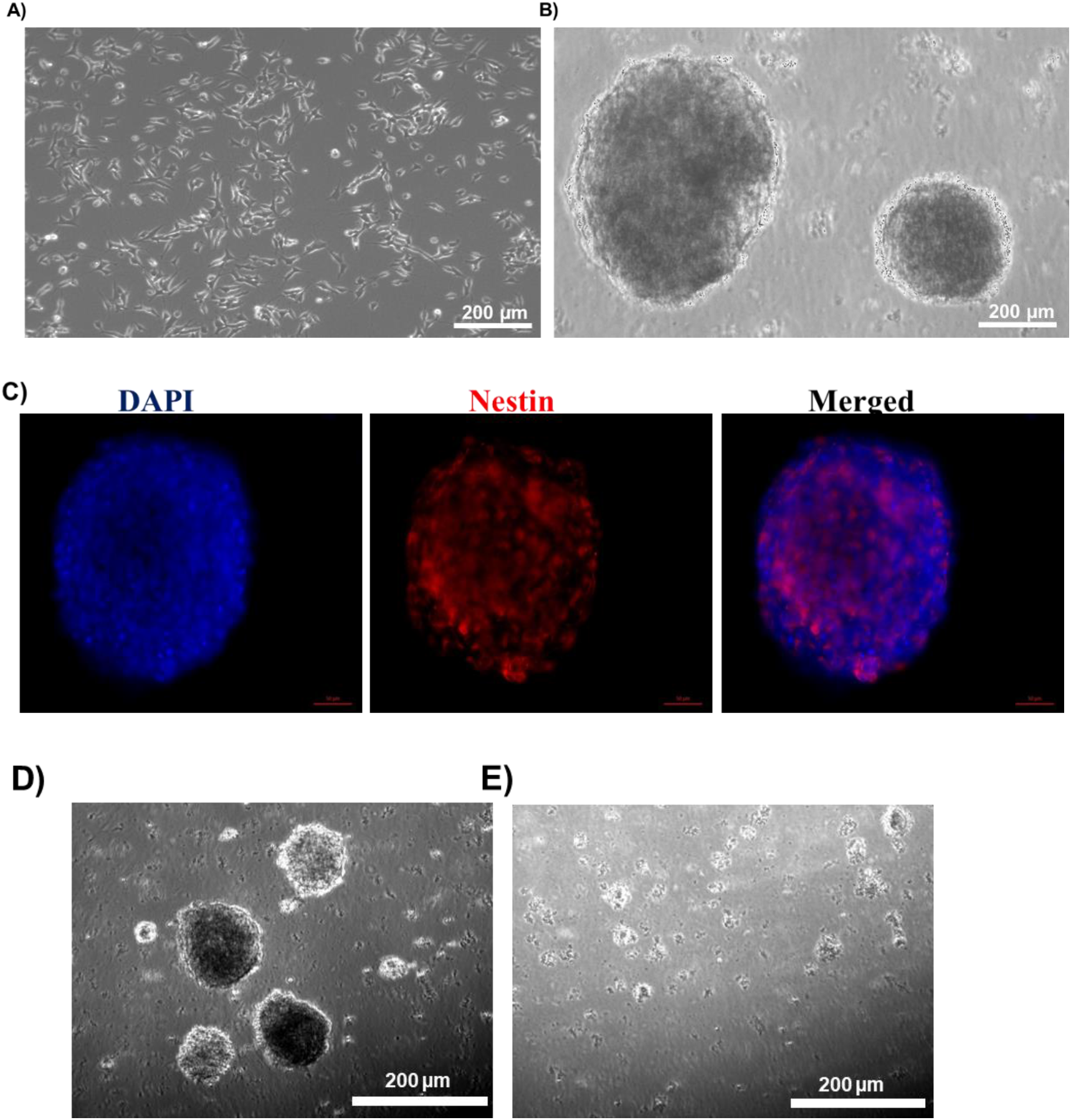
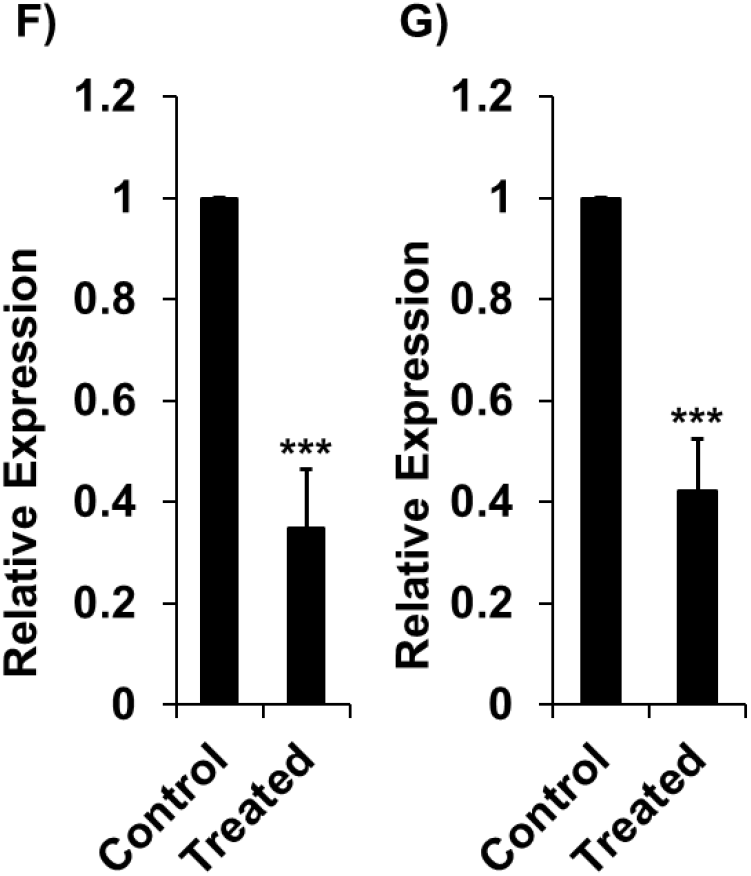
DKK1 treatment reduces the markers of neuroblastoma stem cells proliferation. A) Bright-field image of undifferentiated adherent SHSY5Y cells; B) Bright-field image of SHSY5Y neurosphere. C) Immunocytochemistry images of SHSY5Y neurosphere showing stem cell marker nestin expression. Scale Bar (a and b) 100 μm, Scale Bar (c) 50 μm D) Neurospheres without DKK1 treatment; E) Neurospheres with DKK1 treatment. F-G) Relative mRNA expression of proliferating gene markers Ki67 and PCNA (*** p≤0.0005).

The adherent, undifferentiated SHSY5Y cells tend to grow in clusters and may form clumps, lying one over the other [31]. They show the expression of proliferative gene markers like PCNA and Ki67 [31]. In our study, DKK-1 treatment resulted in a significantly lower expression of proliferation markers (Ki67 and PCNA) in SHSY5Y neurospheres (Figure 1F-G, *** p≤0.0005).

### DKK1 treatment reduces the expression of markers of neuroblastoma stemness and pluripotency

Cancer stem cell surface markers, such as CD133, KIT, and CD44, have been found in many tumors [32, 33]. CD133, KIT, GPRC5C, NOTCH1, PlGF2, TRKB, and LNGFR are the genes that are elevated in neuroblastoma cells as compared to normal stem cells [34]. CD133 and KIT are stem cell markers involved in the proliferation of cells [35]. Overexpression of CD133 is associated with poor clinical outcomes in neuroblastoma and is associated with increased chemoresistance [36]. In our study when we treated SHSY5Y neurospheres to DKK1, we observed a decrease in the expression of stem cell markers. We have found downregulation of cancer stem cell surface markers CD133 and KIT mRNA expression (Figure 2A-B, * p≤0.05, *** p≤0.0005).

**Figure 2:**
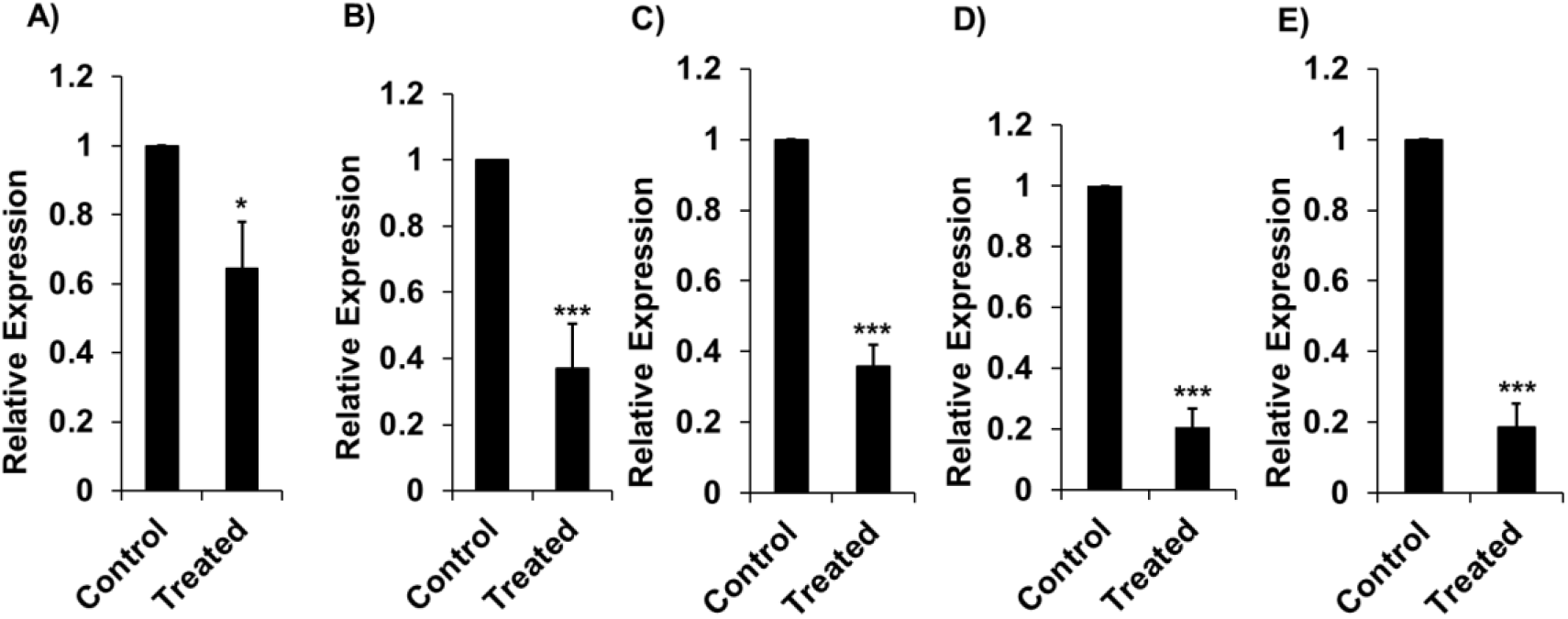
DKK1 treatment reduces the expression of markers of neuroblastoma stemness and pluripotency. (A-B) Relative mRNA expression of cancer stem cell markers CD133 and KIT (*p ≤ 0.05, ***p ≤ 0.0005). (C-E) Relative mRNA expression of pluripotency markers SOX2, OCT4, and NANOG (*** p≤0.0005).

And also, expression of target genes SOX2, OCT4, NANOG is more frequently observed in cancer stem cells than in normal stem cells [37, 38]. SOX2, OCT4, and NANOG are markers of pluripotency; their downregulation results in the perturbation of self-renewable of stem cells. DKK-1 treatment of SHSY5Y neurospheres resulted in significant downregulation of these three pluripotency markers (Figure 2C-E, *** p≤0.0005).

### DKK1 affects β-catenin and TCF mRNA expression

β-catenin is a final effector of the canonical Wnt signaling pathway when affects cell proliferation and differentiation [39, 40]. TCF genes, along with β-catenin act as major transcriptional mediators of Wnt signaling pathways [41]. They determine which genes to be regulated by Wnt signaling. They are responsible for context-dependent interactions of Wnt signaling genes [41]. We studied the effect of DKK1 on β-catenin and TCF gene expression. We found lower mRNA expression of β-catenin, TCF4 and TCF12 after DKK1 treatment (Figure 3A-C, *p ≤ 0.05, ***p ≤ 0.0005).

**Figure 3:**
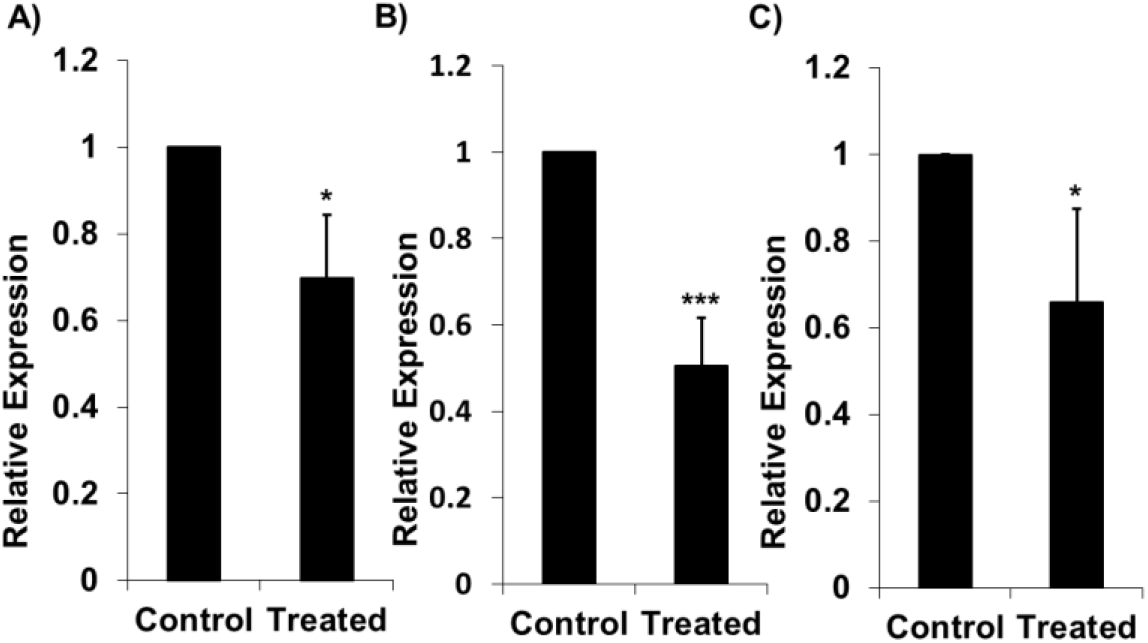
DKK1 affects β-catenin and TCF mRNA expression. DKK1 treatment affects β-catenin and TCF mRNA expression (A-C) Relative mRNA expression of β-catenin, TCF4, and TCF12 (*p ≤ 0.05, ***p ≤ 0.0005).

### DKK1 treatment of adherent SHSY5Y cells induces neuronal differentiation

We studied the effect of DKK1 on SHSY5Y neuronal differentiation. We first treated the SHSY5Y adherent cells in DMEM media with fetal bovine serum with DKK1 only for five days and the control cells were in DMEM media with fetal bovine serum with 0.1 percent BSA in 1X PBS. Morphological differentiation images of SHSY5Y cells without DKK1 and with DKK1 treatment were captured (Figure 4A-B). The relative neurite lengths in samples without DKK1 treatment and in samples with DKK1 treatment were measured (Figure 4C, **p ≤ 0.005). We found significantly higher expression of MAPT (Figure 4D, * p ≤ 0.05), DCX, GAP43, and ENO2 (Figure 4E * p ≤ 0.05) differentiation markers in DKK1 treated cells.

**Figure 4:**
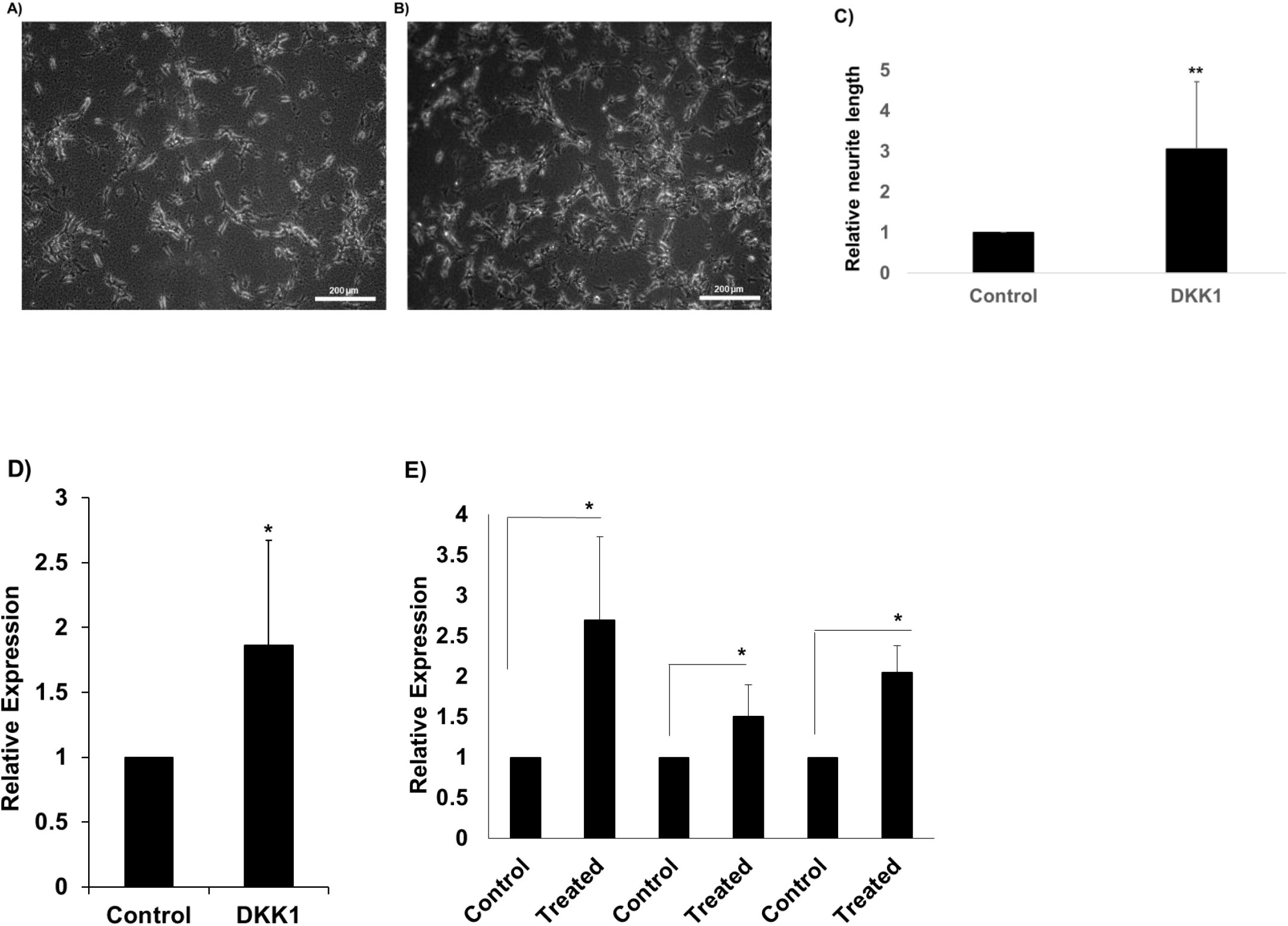
DKK1 treatment of adherent SHSY5Y cells induces neuronal differentiation. DKK1 treatment of adherent SHSY5Y cells induces their neuronal differentiation A-B) Figure showing bright field images of cells without DKK1 treatment and with DKK1 treatment respectively, C) Relative neurite length measured of control sample and DKK1 treated cells (**p ≤ 0.005). D) Relative mRNA expression of MAPT and E) Relative mRNA expression of DCX, GAP43, and ENO2 (*p ≤ 0.05)

### DKK1 synergizes with retinoic acid-induced differentiation of SHSY5Y

In this experiment, we differentiated the adherent SHSY5Y cells using retinoic acid for five days with 0.1 percent BSA in 1X PBS as a control (Figure 5A) and compared this with cells differentiated in media containing retinoic acid and DKK1 for five days (Figure 5B). We have found longer neurites and more differentiated morphology when DKK1 was used with retinoic acid. The relative neurite lengths in samples treated with only retinoic acid and in samples treated with a combination of DKK1 and retinoic acid were measured (Figure 5C, **p ≤ 0.005). We have checked neuronal differentiation markers like GAP43, ENO2, DCX, and MAPT in similar conditions. We have found significantly higher expression of all these differentiation markers in adherent SHSY5Y cells grown in complete DMEM media containing both retinoic acid and DKK1 as compared to cells grown in complete DMEM media containing only retinoic acid (Figure 5D-G, ** p ≤ 0.005, * ≤ 0.05).

**Figure 5:**
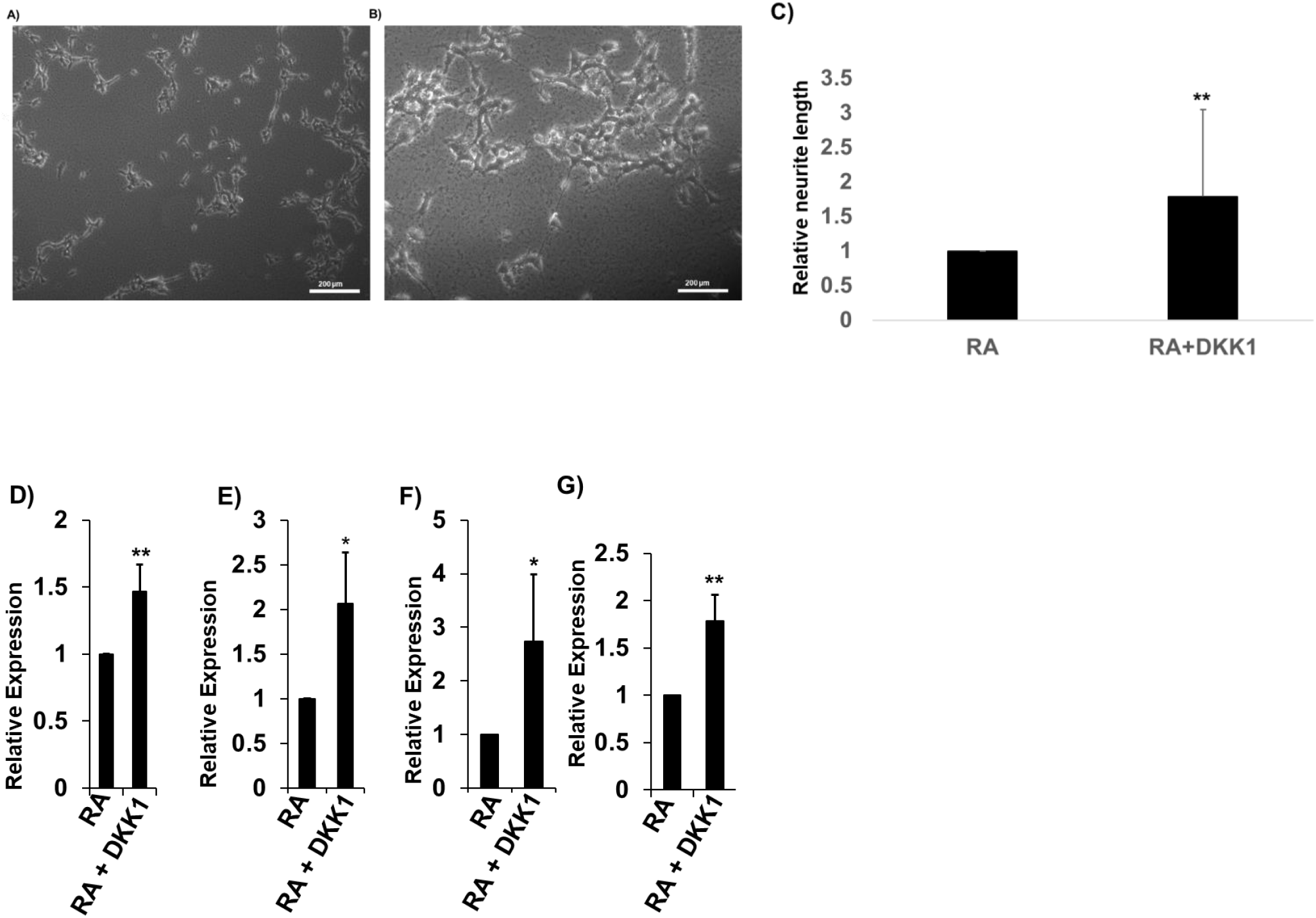
DKK1 synergizes with retinoic acid-induced differentiation of SHSY5Y. DKK1 synergizes with retinoic acid-induced differentiation of SHSY5Y A) Bright field image of adherent day 5 differentiated SHSY5Y treated with retinoic acid only and; B) Bright field image of adherent day 5 differentiated SHSY5Y treated with retinoic acid and DKK1. C) Relative neurite length measured of retinoic acid treated cells and cells treated with combined retinoic acid and DKK1 (**p ≤ 0.005), D-G) Relative mRNA expression of neuronal differentiation markers GAP43, ENO2, DCX, and MAPT (*p ≤ 0.05, **p ≤ 0.005).

## Discussion

Neuroblastoma is a childhood cancer that is developmental in origin, arising from neural progenitor cells outside the nervous system, often from the adrenal region. It is the third-most common cause of child mortality among pediatric cancers. Early onset and detection in a child of up to 0–1-year results in partial or complete cure, but the outlook is less favorable. Different pathways of cellular machinery have been targeted for therapy, but side effects and the development of resistance result in treatment failure. Alk and N-Myc amplification are associated with a poor prognosis. The role of the Wnt pathway and Wnt signaling inhibitors is often context-dependent. This is evident at the level of primary tumors as well as in cell lines.

Several neuroblastoma cell lines have been extensively studied. We worked with SHSY5Y cell line. These lines are grown as adherent cultures, but on the one hand, they can be differentiated into neurons, and on the other, they can form neurospheres with stem cell-like properties under appropriate culture conditions. Considering the role of Wnt signaling in neuroblastoma, we have studied the effects of a Wnt pathway inhibitor, DKK1, on these cells in terms of their stemness-related properties as well as proliferation and differentiation, either by itself or in combination with retinoic acid. DKK1 levels in neuroblastoma are downregulated by MYCN [42], which further supports the logic of supplementing DKK1 levels.

DKK1 can convincingly cause the disintegration of preformed neurospheres of SHSY5Y cells. It is able to lower the expression levels of cancer stem cell markers like CD133 and KIT and pluripotency markers like SOX2, OCT4, and NANOG in SHSY5Y neurospheres. In addition, proliferation markers like Ki67 and PCNA are lowered. This is a novel finding that indicates that a Wnt inhibitor could have a major role in countering the stem-like properties of neuroblastomas. Retinoic acid derivatives are also in clinical use during the maintenance phase of neuroblastoma therapy to prevent relapse after treatment. They have been shown to promote the neuronal differentiation of neuroblastoma-like cells. However, there are limitations to the use of retinoic acid and its derivatives, notably because of the retention of stem cell-like properties in neuroblastoma cells. Considering the abrogation of stem cell-like properties by DKK1, it was then considered if DKK1 could synergize with retinoic acid for inducing cell differentiation. Our results, based on morphological studies including neurite formation and the expression of markers, show that DKK1, by itself, is sufficient for neuronal differentiation in SHSY5Y culture. DKK1 also synergizes with retinoic acid to cause SHSY5Y neuronal differentiation. DKK1 may be considered a candidate molecule by itself or as a prototype molecule for its particular pathway for the inhibition of wnt signaling. The combination of stemness reduction and synergism with retinoic acid for neuronal differentiation makes this molecule or pathway a promising candidate for neuroblastoma management.

## Statements and declarations

### Author Contributions

All authors contributed to the study’s generality and design. Shubham Krishna and Subrata Sinha conceived the idea and planned the experiments. Data collection was done by Shubham Krishna and analysis was performed by Shubham Krishna and Bharat Prajapati. The manuscript was written by Shubham Krishna and Subrata Sinha. All authors read and approved the final handwriting. Pankaj Seth co-supervised the project. Subrata Sinha supervised the project.

## Acknowledgments

We thank Dr. Pankaj Seth’s laboratory for providing us with the SHSY5Y cell line. We also thank our lab technician, P Manish, for his technical assistance. We express our sincere gratitude to the NBRC core fund and the Dr. Subrata Sinha JC Bose fellowship for supporting this project.

## REFERENCES

1. Brodeur GM: Neuroblastoma: biological insights into a clinical enigma. Nat Rev Cancer 2003, 3:203–216.

2. Spix C, Pastore G, Sankila R, Stiller CA, Steliarova-Foucher E: Neuroblastoma incidence and survival in European children (1978-1997): the Automated Childhood Cancer Information System project report. Eur J Cancer 2006, 42:2081–2091.

3. Smith MA, Seibel NL, Altekruse SF, Ries LA, Melbert DL, O’Leary M, Smith FO, Reaman GH: Outcomes for children and adolescents with cancer: challenges for the twenty-first century. J Clin Oncol 2010, 28:2625–2634.

4. Otte J, Dyberg C, Pepich A, Johnsen JI. MYCN Function in Neuroblastoma Development. Front Oncol. 2021 Jan 27;10:624079. doi: 10.3389/fonc.2020.624079. PMID: 33585251; PMCID: PMC7873735.

5. Barbieri E, Mehta P, Chen Z, Zhang L, Slack A, Berg S, Shohet JM: MDM2 inhibition sensitizes neuroblastoma to chemotherapy-induced apoptotic cell death. Mol Cancer Ther 2006, 5:2358–2365.

6. Delbridge AR, Strasser A: The BCL-2 protein family, BH3-mimetics and cancer therapy. Cell Death Differ 2015, 22:1071–1080.

7. Hirsch HA, Iliopoulos D, Tsichlis PN, Struhl K: Metformin selectively targets cancer stem cells, and acts together with chemotherapy to block tumor growth and prolong remission. Cancer Res 2009, 69:7507–7511.

8. Moreira PI, Custodio J, Moreno A, Oliveira CR, Santos MS: Tamoxifen and estradiol interact with the flavin mononucleotide site of complex I leading to mitochondrial failure. J Biol Chem 2006, 281:10143–10152.

9. Berardi DE, Campodonico PB, Diaz Bessone MI, Urtreger AJ, Todaro LB: Autophagy: friend or foe in breast cancer development, progression, and treatment. Int J Breast Cancer 2011, 2011:595092.

10. Wang SY, Yu QJ, Zhang RD, Liu B: Core signaling pathways of survival/death in autophagy-related cancer networks. Int J Biochem Cell Biol 2011, 43:1263–1266.

11. Izycka-Swieszewska E, Drozynska E, Rzepko R, Kobierska-Gulida G, Grajkowska W, Perek D, Balcerska A: Analysis of PI3K/AKT/mTOR signalling pathway in high-risk neuroblastic tumours. Pol J Pathol 2010, 61:192–198.

12. Biedler JL, Roffler-Tarlov S, Schachner M, Freedman LS. Multiple Neurotransmitter Synthesis by Human Neuroblastoma Cell Lines and Clones. Cancer Research. 1978;38(11 Pt 1):3751–3757.

13. Kovalevich J, Santerre M, Langford D. Considerations for the Use of SH-SY5Y Neuroblastoma Cells in Neurobiology. Methods Mol Biol. 2021;2311:9–23. doi: 10.1007/978-1-0716-1437-2_2. PMID: 34033074.

14. Påhlman S, Hoehner JC, et al. Differentiation and survival influences of growth factors in human neuroblastoma. European Journal of Cancer. 1995;31(4):453–458.

15. Matthay KK, Reynolds CP, Seeger RC, Shimada H, Adkins ES, Haas-Kogan D, Gerbing RB, London WB, Villablanca JG: Long-term results for children with high-risk neuroblastoma treated on a randomized trial of myeloablative therapy followed by 13-cis-retinoic acid: a children’s oncology group study. J Clin Oncol 2009, 27:1007–1013.

16. Brodeur GM, Seeger RC, Schwab M, Varmus HE, Bishop JM. Amplification of N-myc in untreated human neuroblastomas correlates with advanced disease stage. Science. 1984 Jun 8;224(4653):1121–4. doi: 10.1126/science.6719137. PMID: 6719137.

17. Duffy DJ, Krstic A, Halasz M, Schwarzl T, Konietzny A, Iljin K, Higgins DG, Kolch W. Retinoic acid and TGF-β signalling cooperate to overcome MYCN-induced retinoid resistance. Genome Med. 2017 Feb 10;9(1):15. doi: 10.1186/s13073-017-0407-3. PMID: 28187790; PMCID: PMC5303304.

18. Bayeva N, Coll E, Piskareva O. Differentiating Neuroblastoma: A Systematic Review of the Retinoic Acid, Its Derivatives, and Synergistic Interactions. J Pers Med. 2021 Mar 16;11(3):211. doi: 10.3390/jpm11030211. PMID: 33809565; PMCID: PMC7999600.

19. Zhan T, Rindtorff N, Boutros M. Wnt signaling in cancer. Oncogene. 2017 Mar;36(11):1461–1473. doi: 10.1038/onc.2016.304. Epub 2016 Sep 12. PMID: 27617575; PMCID: PMC5357762.

20. Zhang J, Zhou B, Liu Y, Chen K, Bao P, Wang Y, Wang J, Zhou Z, Sun X, Li Y. Wnt inhibitory factor-1 functions as a tumor suppressor through modulating Wnt/β-catenin signaling in neuroblastoma. Cancer Lett. 2014 Jun 28;348(1-2):12–9. doi: 10.1016/j.canlet.2014.02.011. Epub 2014 Feb 18. PMID: 24561119.

21. Szemes M, Greenhough A, Melegh Z, Malik S, Yuksel A, Catchpoole D, Gallacher K, Kollareddy M, Park JH, Malik K. Wnt Signalling Drives Context-Dependent Differentiation or Proliferation in Neuroblastoma. Neoplasia. 2018 Apr;20(4):335–350. doi: 10.1016/j.neo.2018.01.009. Epub 2018 Mar 3. PMID: 29505958; PMCID: PMC5909736.

22. Becker J, Wilting J. WNT signaling, the development of the sympathoadrenal-paraganglionic system and neuroblastoma. Cell Mol Life Sci. 2018 Mar;75(6):1057–1070. doi: 10.1007/s00018-017-2685-8. Epub 2017 Oct 22. PMID: 29058015; PMCID: PMC5814469.

23. Becker J, Wilting J. WNT Signaling in Neuroblastoma. Cancers (Basel). 2019 Jul 19;11(7):1013. doi: 10.3390/cancers11071013. PMID: 31331081; PMCID: PMC6679057.

24. Suebsoonthron J, Jaroonwitchawan T, Yamabhai M, Noisa P. Inhibition of WNT signaling reduces differentiation and induces sensitivity to doxorubicin in human malignant neuroblastoma SH-SY5Y cells. Anticancer Drugs. 2017 Jun;28(5):469–479. doi: 10.1097/CAD.0000000000000478. PMID: 28240680.

25. Tang Q, Chen J, Di Z, Yuan W, Zhou Z, Liu Z, Han S, Liu Y, Ying G, Shu X, Di M. TM4SF1 promotes EMT and cancer stemness via the Wnt/β-catenin/SOX2 pathway in colorectal cancer. J Exp Clin Cancer Res. 2020 Nov 5;39(1):232. doi: 10.1186/s13046-020-01690-z. PMID: 33153498; PMCID: PMC7643364.

26. Cho YH, Ro EJ, Yoon JS, Mizutani T, Kang DW, Park JC, Il Kim T, Clevers H, Choi KY. 5-FU promotes stemness of colorectal cancer via p53-mediated WNT/β-catenin pathway activation. Nat Commun. 2020 Oct 21;11(1):5321. doi: 10.1038/s41467-020-19173-2. PMID: 33087710; PMCID: PMC7578039.

27. Chen J, Wang P, Cai R, Peng H, Zhang C, Zhang M. SLC34A2 promotes neuroblastoma cell stemness via enhancement of miR-25/Gsk3β-mediated activation of Wnt/β-catenin signaling. FEBS Open Bio. 2019 Feb 7;9(3):527–537. doi: 10.1002/2211-5463.12594. PMID: 30868061; PMCID: PMC6396163.

28. Flahaut M, Meier R, Coulon A, Nardou KA, Niggli FK, Martinet D, Beckmann JS, Joseph JM, Mühlethaler-Mottet A, Gross N. The Wnt receptor FZD1 mediates chemoresistance in neuroblastoma through activation of the Wnt/beta-catenin pathway. Oncogene. 2009 Jun 11;28(23):2245–56. doi: 10.1038/onc.2009.80. Epub 2009 May 4. PMID: 19421142.

29. Szemes M, Greenhough A, Melegh Z, Malik S, Yuksel A, Catchpoole D, Gallacher K, Kollareddy M, Park JH, Malik K. Wnt Signalling Drives Context-Dependent Differentiation or Proliferation in Neuroblastoma. Neoplasia. 2018 Apr;20(4):335–350. doi: 10.1016/j.neo.2018.01.009. Epub 2018 Mar 3. PMID: 29505958; PMCID: PMC5909736.

30. Zhang, Jiao et al. “Expression and clinical significance of Dickkofp-1 (DKK-1) in neuroblastoma.” (2017).

31. Kovalevich J, Santerre M, Langford D. Considerations for the Use of SH-SY5Y Neuroblastoma Cells in Neurobiology. Methods Mol Biol. 2021;2311:9–23. doi: 10.1007/978-1-0716-1437-2_2. PMID: 34033074.

32. Shimokawa M, Ohta Y, Nishikori S, Matano M, Takano A, Fujii M, Date S, Sugimoto S, Kanai T, Sato T. Visualization and targeting of LGR5<sup>+</sup> human colon cancer stem cells. Nature. 2017 May 11;545(7653):187–192. doi: 10.1038/nature22081. Epub 2017 Mar 29. PMID: 28355176.

33. Shibata M, Hoque MO. Targeting Cancer Stem Cells: A Strategy for Effective Eradication of Cancer. Cancers (Basel). 2019 May 27;11(5):732. doi: 10.3390/cancers11050732. PMID: 31137841; PMCID: PMC6562442.

34. Ross RA, Walton JD, Han D, Guo HF, Cheung NK. A distinct gene expression signature characterizes human neuroblastoma cancer stem cells. Stem Cell Res. 2015 Sep;15(2):419–26. doi: 10.1016/j.scr.2015.08.008. Epub 2015 Aug 20. PMID: 26342562; PMCID: PMC4601571.

35. Singh SK, Hawkins C, Clarke ID, Squire JA, Bayani J, Hide T, Henkelman RM, Cusimano MD, Dirks PB. Identification of human brain tumour initiating cells. Nature. 2004 Nov 18;432(7015):396–401. doi: 10.1038/nature03128. PMID: 15549107.

36. Tong QS, Zheng LD, Tang ST, Ruan QL, Liu Y, Li SW, Jiang GS, Cai JB. Expression and clinical significance of stem cell marker CD133 in human neuroblastoma. World J Pediatr. 2008 Feb;4(1):58–62. doi: 10.1007/s12519-008-0012-z. PMID: 18402255.

37. Basati G, Mohammadpour H, Emami Razavi A. Association of High Expression Levels of SOX2, NANOG, and OCT4 in Gastric Cancer Tumor Tissues with Progression and Poor Prognosis. J Gastrointest Cancer. 2020 Mar;51(1):41–47. doi: 10.1007/s12029-018-00200-x. PMID: 30628031.

38. Wang X, Liu Q, Hou B, Zhang W, Yan M, Jia H, Li H, Yan D, Zheng F, Ding W, Yi C, Hai Wang. Concomitant targeting of multiple key transcription factors effectively disrupts cancer stem cells enriched in side population of human pancreatic cancer cells. PLoS One. 2013 Sep 11;8(9):e73942. doi: 10.1371/journal.pone.0073942. PMID: 24040121; PMCID: PMC3770686.

39. Becker J, Wilting J. WNT signaling, the development of the sympathoadrenal-paraganglionic system and neuroblastoma. Cell Mol Life Sci. 2018 Mar;75(6):1057–1070. doi: 10.1007/s00018-017-2685-8. Epub 2017 Oct 22. PMID: 29058015; PMCID: PMC5814469.

40. Becker J, Wilting J. WNT Signaling in Neuroblastoma. Cancers (Basel). 2019 Jul 19;11(7):1013. doi: 10.3390/cancers11071013. PMID: 31331081; PMCID: PMC6679057.

41. Pai SG, Carneiro BA, Mota JM, Costa R, Leite CA, Barroso-Sousa R, Kaplan JB, Chae YK, Giles FJ. Wnt/beta-catenin pathway: modulating anticancer immune response. J Hematol Oncol. 2017 May 5;10(1):101. doi: 10.1186/s13045-017-0471-6. PMID: 28476164; PMCID: PMC5420131.

42. Koppen A, Ait-Aissa R, Hopman S, Koster J, Haneveld F, Versteeg R, Valentijn LJ. Dickkopf-1 is down-regulated by MYCN and inhibits neuroblastoma cell proliferation. Cancer Lett. 2007 Oct 28;256(2):218–28. doi: 10.1016/j.canlet.2007.06.011. Epub 2007 Jul 23. PMID: 17643814.

