## Supplementary Figure S1 for "DKK1, a negative regulator of Wnt signaling, is a novel inducer of neuroblastoma differentiation"

| GENE NAME | FORWARD PRIMER SEQUENCE | REVERSE PRIMER SEQUENCE |
| --- | --- | --- |
| CD133 | TTTGAAACAGAGGAAGCAGAGA | GCATTTAGACATCTTCCTAGGGGT |
| KIT | CACCGAAGGAGGCACTTACA | CAAACACGAGCCACAACTTT |
| SOX2 | ACACCAATCCCATCCACACT | GCAAACTTCCTGCAAAGCTC |
| OCT4 | GTAGGGTAAAGGAGGGAAGGAGA | ATCTACCTAGCCACCAGACCA |
| NANOG | TAGCAATGGTGTGACGCAGG | TGTCTGTGACTGGAGTTGTGT |
| Ki67 | GCCTGCTCGACCCTACAGA | GCTTGTCAACTGCGGTTGC |
| PCNA | GCGTGAACCTCACCAGTATGT | TCTTCGGCCCTTAGTGTAATGAT |
| BETA CATENIN | TGCAGTTCGCCTTCACTATGGACT | GATTTGCGGGACAAAGGGCAAGAT |
| TCF4 | TGCAAAGCCGAATTGAAGATCG | AGAAGGTCCAATGATTCCATGC |
| TCF12 | CCAGTAGTTATGGCAACCTTCAT | GACTCGTGTTTATGTCTGTTGGT |
| MAPT | CCAAGTGTGGCTCATTAGGCA | CCAATCTTCGACTGGACTCTGT |
| DCX | AATCACCAAGCGAGTCCGAG | AAAGCAGACATTCCAGAGCTCAA |
| GAP43 | GGCCGCAACCAAAATTCAGG | CGGCAGTAGTGGTGCCTTC |
| ENO2 | AGGTGCAGAGGTCTACCATAC | AGCTCCAAGGCTTCACTGTTC |
| GAPDH | ACATCAAGAAGGTGGTGAAGCAGG | TGTCGCTGTTGAAGTCAGAGGAGA |
